# Directing voluntary temporal attention increases fixational stability

**DOI:** 10.1101/449264

**Authors:** Rachel N. Denison, Shlomit Yuval-Greenberg, Marisa Carrasco

**Affiliations:** Department of Psychology and Center for Neural Science, New York University, New York, NY 10003, USA; School of Psychological Sciences and Sagol School of Neuroscience, Tel Aviv University, Tel Aviv, 6997801, Israel

## Abstract

Our visual input is constantly changing, but not all moments are equally relevant. Temporal attention, the prioritization of visual information at specific points in time, increases perceptual sensitivity at behaviorally relevant times. The dynamic processes underlying this increase are unclear. During fixation, humans make small eye movements called microsaccades, and inhibiting microsaccades improves perception of brief stimuli. Here we asked whether temporal attention changes the pattern of microsaccades in anticipation of brief stimuli. Human observers (female and male) judged brief stimuli presented within a short sequence. They were given either an informative precue to attend to one of the stimuli, which was likely to be probed, or an uninformative (neutral) precue. We found strong microsaccadic inhibition before the stimulus sequence, likely due to its predictable onset. Critically, this anticipatory inhibition was stronger when the first target in the sequence (T1) was precued (task-relevant) than when the precue was uninformative. Moreover, the timing of the last microsaccade before T1 and the first microsaccade after T1 shifted, such that both occurred earlier when T1 was precued than when the precue was uninformative. Finally, the timing of the nearest pre- and post-T1 microsaccades affected task performance. Directing voluntary temporal attention therefore impacts microsaccades, helping to stabilize fixation at the most relevant moments, over and above the effect of predictability. Just as saccading to a relevant stimulus can be an overt correlate of the allocation of spatial attention, precisely timed gaze stabilization can be an overt correlate of the allocation of temporal attention.

**Significance statement:** We pay attention at moments in time when a relevant event is likely to occur. Such temporal attention improves our visual perception, but how it does so is not well understood. Here we discovered a new behavioral correlate of voluntary, or goal-directed, temporal attention. We found that the pattern of small fixational eye movements called microsaccades changes around behaviorally relevant moments in a way that stabilizes the position of the eyes. Microsaccades during a brief visual stimulus can impair perception of that stimulus. Therefore, such fixation stabilization may contribute to the improvement of visual perception at attended times. This link suggests that in addition to cortical areas, subcortical areas mediating eye movements may be recruited with temporal attention.

## Introduction

Temporal attention is the prioritization of sensory information at specific points in time. It allows us to combine information about the expected timing of sensory events with ongoing task goals to improve our perception and behavior (Nobre and Rohenkohl, 2014; Nobre and van Ede, 2018). For example, when returning a tennis serve, it is critical to see the ball well at the moment it meets your opponent’s racket but less critical to see it well a half second before. Voluntarily directing attention to a relevant time increases perceptual sensitivity at that time (Correa et al., 2006; Davranche et al., 2011; Rohenkohl et al., 2014; Samaha et al., 2015; Denison et al., 2017) and decreases sensitivity at other times, resulting in attentional tradeoffs (Denison et al., 2017). The mechanisms underlying the allocation and perceptual effects of voluntary temporal attention remain poorly understood.

To understand attention as a dynamic process, it is critical to distinguish between temporal attention-the prioritization of a task–relevant time–and temporal expectation–the ability to predict stimulus timing, regardless of task-relevance. The conceptual distinction between attention (relevance) and expectation (predictability) has been established in the spatial and feature domains, in which the two factors have dissociable impacts on perception and neural responses (Summerfield and Egner, 2009; Kok et al., 2012; Wyart et al., 2012; Summerfield and Egner, 2016). Here we manipulated temporal attention while equating expectation by using precues to direct voluntary temporal attention to specific stimuli in predictably timed sequences of brief visual targets (Denison et al., 2017). This task requires temporally precise cognitive control to attend to a relevant time point that varies from trial to trial.

We investigated the possibility of an overt, oculomotor signature of voluntary temporal attention by examining the interaction between voluntary temporal attention and microsaccades. Microsaccades are small (<1°) eye movements made 1-2 times per second even while fixating (Otero-Millan et al., 2008; Rolfs, 2009; Martinez-Conde et al., 2013; Rucci and Poletti, 2015). Correlations have been observed between microsaccades and *spatial* covert attention during fixation (Hafed and Clark, 2002; Engbert and Kliegl, 2003; Yuval-Greenberg et al., 2014; Lowet et al., 2018). However, their interpretation has been controversial (Tse et al., 2004; Martinez-Conde et al., 2013), and spatial attention also affects behavior in the absence of microsaccades (Poletti et al., 2017). Importantly for understanding the mechanisms underlying temporal attention, microsaccades provide a continuous, online physiological measure that can reveal dynamic processes unavailable from behavioral measures alone.

Microsaccades contribute to variability in visual perception (Hafed et al., 2015). They enhance perception of static stimuli by preventing and counteracting perceptual fading during sustained fixation (Ditchburn and Ginsborg, 1952; Riggs et al., 1953; Martinez-Conde et al., 2006; McCamy et al., 2012; Costela et al., 2017) and can correspondingly increase neural activity (Martinez-Conde et al., 2013; Troncoso et al., 2015). They also improve vision by positioning stimuli at the highest-acuity region within the fovea (Poletti et al., 2013; Rucci and Poletti, 2015). However, microsaccades impair perception of brief stimuli that occur around the time of a microsaccade onset, a phenomenon known as microsaccadic suppression (Zuber and Stark, 1966; Beeler, 1967; Hafed and Krauzlis, 2010; Hafed et al., 2011; Amit et al., in press). Corresponding neural response reductions just after microsaccades have been found in multiple visual areas (Herrington et al., 2009; Hafed and Krauzlis, 2010; Martinez-Conde et al., 2013; Chen et al., 2015; Chen and Hafed, 2017; Loughnane et al., 2018).

Humans and monkeys actively inhibit microsaccades before a predictably timed, brief stimulus, which helps avoid microsaccadic suppression during the stimulus presentation (Findlay, 1974; Betta and Turatto, 2006; Pastukhov and Braun, 2010; Hafed et al., 2011; Fried et al., 2014; Dankner et al., 2017; Olmos-Solis et al., 2017; Amit et al., in press). Recent studies have linked temporal predictability, pre-target microsaccade inhibition and performance improvement. Specifically, neurotypical adults inhibit microsaccades more before predictably timed stimuli than randomly timed stimuli (Dankner et al., 2017; Amit et al., in press), whereas adults with ADHD fail to do so (Dankner et al., 2017). Therefore, the control of microsaccade timing in accordance with temporal expectations improves performance, and may play a role in performance impairments in clinical populations.

Whether microsaccade dynamics are also sensitive to temporal attention is unknown. It has not been investigated whether microsaccades are controlled to prioritize more relevant over less relevant stimulus times, when all stimuli are equally predictable. Here we manipulated temporal attention using a precue and examined the effect of this manipulation on microsaccades. We found that, beyond the effects of expectation, directing temporal attention increases the stabilization of eye position at the time of a brief, relevant visual stimulus.

## Methods

### Data set

We reanalyzed eye-tracking data collected in a recent study on temporal attention by Denison, Heeger and Carrasco (2017). Thus behavioral procedures were identical to those previously reported. To maximize power of the microsaccade analysis, we combined the data from all three experiments in that study. The stimuli and tasks were similar across experiments: on each trial, human observers were presented with a predictably timed sequence of two or three target gratings–which we refer to as T1, T2 and T3–and judged the orientation of one of these gratings. Precues before each sequence directed temporal attention to one or more grating times. The experiments varied as follows: Experiment 1 used an orientation discrimination task with 2-target sequences. Experiment 2 used an orientation discrimination task with 3-target sequences. Experiment 3 used an orientation estimation task with 2-target sequences. All grating stimuli were potential targets and we refer to them as such. To combine data across experiments, we focused on the precue conditions that were common to all experiments (neutral, precue T1, precue T2; see “Behavioral procedures”). We also analyzed precue T3 trials from Experiment 2 when appropriate.

### Observers

The observers were the same as in Denison et al. (2017), except that eye-tracking data from five observers (three in Experiment 1, one in Experiments 2 and 3) could not be used for microsaccade analysis for technical reasons (e.g., insufficient sampling rate). To better equate the number of observers in each experiment for the present study, we collected data from three new observers for Experiment 1. This gave 30 total data sets: 10 in Experiment 1, 9 in Experiment 2, and 11 in Experiment 3. Four observers participated in multiple experiments: one observer participated in Experiments 1 and 2; two observers participated in Experiments 1 and 3; and one observer (R.N.D) participated in all three experiments. Therefore, 25 unique observers (16 female, 9 male) are included in the present study. All observers provided informed consent, and the University Committee on Activities involving Human Subjects at New York University approved the experimental protocols. All observers had normal or corrected-to-normal vision.

### Stimuli

Stimuli were generated on an Apple iMac using Matlab and Psychophysics Toolbox (Brainard, 1997; Pelli, 1997; Kleiner et al., 2007). They were displayed on a gamma-corrected Sony Trinitron G520 CRT monitor with a refresh rate of 100 Hz at a viewing distance of 56 cm. Observers’ heads were stabilized by a head rest. A central white fixation “x” subtended 0.5° visual angle. Visual target stimuli were 4 cpd sinusoidal gratings with a 2D Gaussian spatial envelope (standard deviation 0.7°), presented in the lower right quadrant of the display centered at 5.7° eccentricity (**Figure 1a**). Stimuli were high contrast (64% or 100%, which we combined as there were no behavioral differences). Placeholders, corners of a 4.25° x 4.25° white square outline (line width 0.08°) centered on the target location, were present throughout the display to minimize spatial uncertainty. The stimuli were presented on a medium gray background (57 cd/m^2^). In Experiments 1 and 3, in which there were two target stimuli, auditory precues were high (precue T1: 784 Hz; G5) or low (precue T2: 523 Hz; C5) pure sine wave tones, or their combination (neutral precue). In Experiment 2, in which there were three target stimuli, auditory precues were high (precue T1: 1318 Hz; E6), medium (precue T2: 784 Hz; G5), or low (precue T3: 330 Hz; E4) tones, or their combination (neutral precue). Auditory stimuli were presented on the computer speakers.

**Figure 1.**
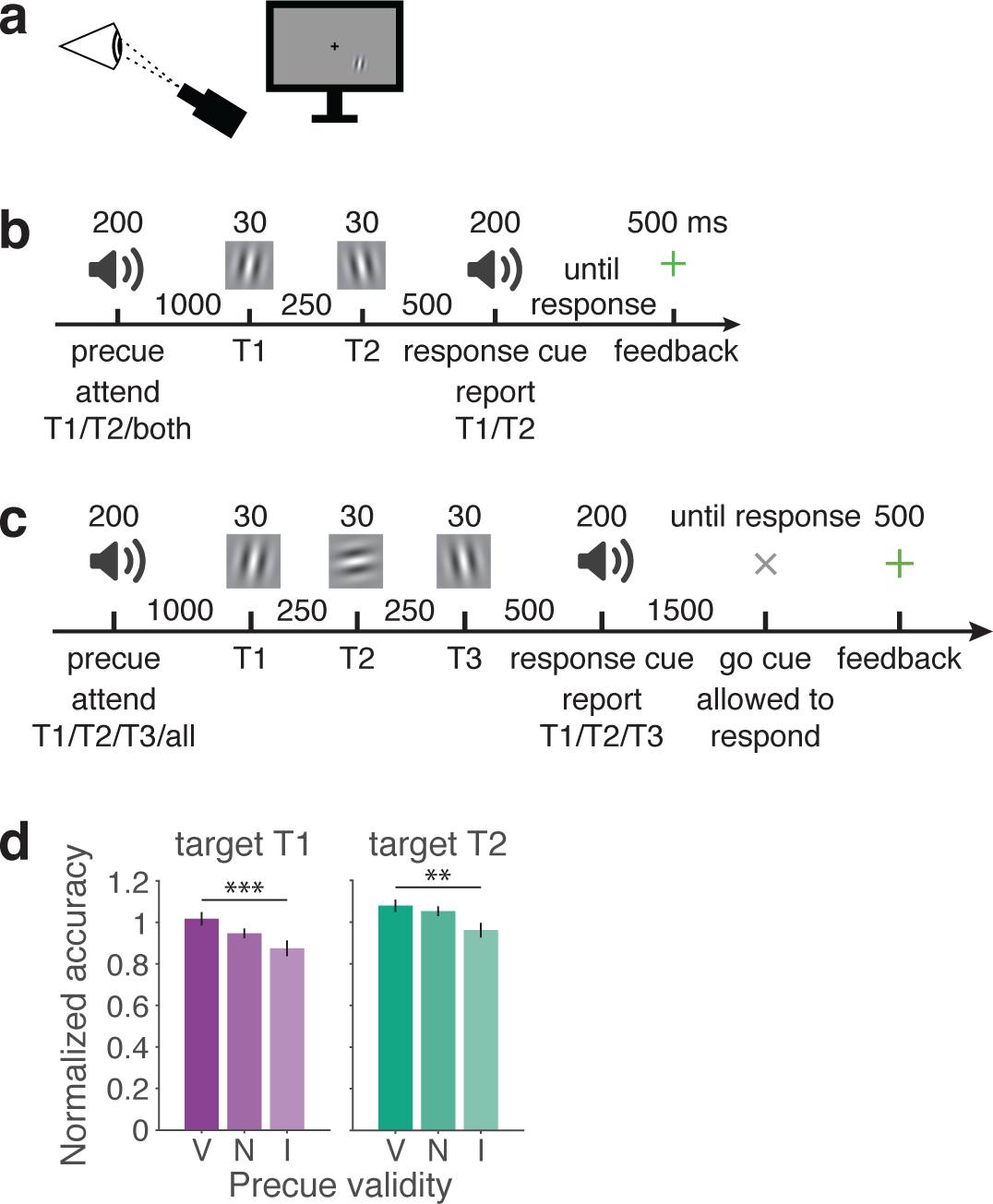
Task and behavior. **a)** Schematic of eye tracking and display setup. Observers fixated on a central cross, and all stimuli appeared in the lower right quadrant. **b)** Trial timeline for 2-target tasks (Experiments 1 and 3). In Experiment 3, a probe grating appeared after the response cue, which the observer adjusted to estimate orientation (not shown). **c)** Trial timeline for 3-target task (Experiment 2). **d)** Performance accuracy normalized to average neutral performance for each observer, mean and SEM. Experiments 1-3, n=30. V=valid, N=neutral, I=invalid. **p<0.01, ***p<0.001.

### Experimental design and statistical analysis

Thirty data sets were analyzed (see *Observers*). The sample size was determined by Denison et al. (2017), as we reanalyzed the data collected in that study. The within-observers factors were precue type and/or target. The between-observers factor was experiment.

Correction for multiple comparisons was achieved using non-parametric methods or the Bonferroni method. The Methods subsection *Data analysis* contains “Statistics” subsections for each analysis, which describe all statistical procedures. Statistical analyses were performed using *R*.

### Behavioral procedures

#### Basic task and trial sequence

Observers judged the orientation of grating patches that appeared in short sequences of two or three target stimuli per trial (Experiments 1 and 3: two targets, T1, T2; Experiment 2: three targets, T1, T2, T3). Targets were presented for 30 ms each at the same spatial location, separated by stimulus onset asynchronies (SOAs) of 250 ms (**Figure 1b,c**). An auditory precue 1000 ms before the first target instructed observers to attend to one of the targets (informative precue, single tone) or to sustain attention across all targets (neutral precue, all tones simultaneously). Observers were asked to report the orientation of one of the targets, which was indicated by an auditory response cue 500 ms after the last target (same tones as informative precues). The duration of the precue and response cue tones was 200 ms. The timing of auditory and visual events was the same on every trial. From trial to trial, the allocation of temporal attention varied (depending on the precue), and the response selection varied (depending on the response cue).

#### Attention manipulation

On valid trials (60% of trials), the response cue matched the precue; observers were asked to report the same target they had been instructed to attend. On invalid trials (20%), the response cue mismatched the precue; observers were asked to report a different target than the one they had been instructed to attend. Thus informative precues were 75% valid. On neutral trials (20%), observers were given a neutral, uninformative precue, and they were equally likely to be asked to report any of the targets. Thus observers had incentive to attend to the target indicated by the precue, because they were most likely to be asked to report its orientation at the end of the trial.

#### Online fixation monitoring

Online streaming of gaze positions was used to ensure central fixation throughout the experiment (see *Eye tracking procedures*). Initiation of each trial was contingent on fixation, with a 750 ms minimum inter-trial interval. Observers were required to maintain fixation, without blinking, from the onset of the precue until the onset of the response cue. If observers broke fixation during this period, the trial was stopped and repeated at the end of the block.

#### Discrimination task

In Experiments 1 and 2, observers performed an orientation discrimination task (**Figure 1b,c**). Each target was tilted slightly clockwise (CW) or counterclockwise (CCW) from either the vertical or horizontal axis, with independent tilts and axes for each target, and observers pressed a key to report the tilt (CW or CCW) of the target indicated by the response cue, with unlimited time to respond. Tilt magnitudes were determined separately for each observer by a thresholding procedure before the main experiment. Observers received feedback at fixation (correct: green “+”; incorrect: red “-”) after each trial, as well as feedback about performance accuracy (percent correct) following each experimental block.

#### Estimation task

In Experiment 3, observers performed an orientation estimation task (**Figure 1b**). Target orientations were selected randomly and uniformly from 0-180°, with independent orientations for each target. Observers estimated the orientation of the target indicated by the response cue by adjusting a grating probe to match the perceived target orientation. The probe was identical to the target but appeared in a new random orientation. Observers moved the mouse horizontally to adjust the orientation of the probe and clicked the mouse to submit the response, with unlimited time to respond. The absolute difference between the reported and presented target orientation was the error for that trial. Observers received feedback at fixation after each trial (error <5°, green “+”; 5-10°, yellow “+”; ≥10°, red “-”). Additional feedback after each block showed the percent of trials with <5° errors, which were defined to observers as “correct”.

#### Training and testing sessions

All observers completed one session of training prior to the experiment to familiarize them with the task and, in Experiments 1 and 2, determine their tilt thresholds. Thresholds were selected to achieve ~79% performance on neutral trials. Observers completed 640 trials across 2 one-hour sessions in Experiments 1 and 3 and 960 trials across 3 sessions in Experiment 2. All experimental conditions were randomly interleaved across trials.

### Eye tracking procedures

#### Eye data collection

Eye position was recorded using an EyeLink 1000 eye tracker (SR Research) with a sampling rate of 1000 Hz. Raw gaze positions were converted into degrees of visual angle using the 5-point-grid calibration, which was performed at the start of each experimental run.

#### Eye data preprocessing

Gaze-position data were segmented into epochs from −1500 to 1250 ms relative to the onset of T1. Blink intervals were identified in these segments according to the EyeLink blink-detection algorithm, along with samples from 200 ms preceding to 200 ms following each blink. (Blinks could occur before the precue or after the response cue.) Blink intervals were removed from the data prior to microsaccade detection. Microsaccades were detected on low-pass filtered data (at 60Hz) using an established algorithm (Engbert and Kliegl, 2003) that compares eye-movement velocity with a threshold criterion set individually for each trial. The threshold was determined on the basis of the 2-D (horizontal and vertical) eye-movement velocity during the trial segment. We set the threshold to be 6 times the standard deviation of the 2-D eye-movement velocity, using a median-based estimate of the standard deviation (Engbert and Kliegl, 2003). A microsaccade was identified when the eye-movement velocity exceeded this threshold for at least 6 ms (seven consecutive eye-position samples). We also imposed a minimum intersaccadic interval (defined as the interval between the last sample of one saccade and the first sample of the next saccade) of 50 ms so that potential overshoot corrections were not considered new microsaccades. We excluded saccades that were larger than 1 degree of visual angle. The time, amplitude, velocity, and direction of each microsaccade were recorded.

### Data analysis

#### Behavioral data analysis

To combine behavioral data across discrimination and estimation experiments, we first calculated accuracy for the estimation experiment (Experiment 3). We assigned accuracy for each trial based on the feedback provided to observers during the experiment, where an estimation report within 5° of the true stimulus orientation was considered “correct”. This accuracy measure was also used for the analysis of behavior vs. microsaccade timing, where a trial-by-trial measure was required. Observer accuracies according to this criterion tended to be lower than for the discrimination experiments (chance performance to be within ±5° of the true orientation was 10°/180° = 5.6% vs. 50% for discrimination). Therefore, to combine data, we first normalized the data from each observer to the neutral condition, dividing the accuracy for each condition (valid, neutral, and invalid for T1 and T2) by the average accuracy across T1 and T2 neutral conditions. So normalized accuracies >1 were better than the observer’s average neutral performance and <1 were worse.

*Statistics*. We used a linear mixed model to evaluate the effects of precue validity and experiment on normalized accuracy in the combined data set, separately for each target. We tested for main effects and interactions by approximating likelihood ratio tests to compare models with and without the effect of interest. Our main interest was confirming the behavioral effect of precue validity on accuracy in the full data set of the 3 experiments combined, which was expected from (Denison et al., 2017), where each experiment was analyzed separately.

#### Microsaccade rate analysis

Microsaccade-rate time-courses were calculated by averaging the number of microsaccade onsets per time sample across all trials, multiplying these values by the sampling rate (1000 Hz) and then smoothing across time by applying a sliding window of 50ms.

Mean microsaccade rate for each precue type was calculated for each observer. Precue types were: precue T1, precue T2, and neutral in all experiments (n=30), and precue T3 in Experiment 2 only, because this was the only experiment with three targets (n=9). Note that, as described in Methods, trials with precues to specific targets could be valid or invalid trials, depending on the response cue. For example, precue T1 trials included T1 valid trials (response cue T1) and T2 invalid trials (response cue T2). The response cue is irrelevant for the present microsaccade analysis, because it occurs at the end of the trial.

*Statistics*. To analyze our data set – a repeated measures design in three separate experiments, with some observers in multiple experiments – we used a linear mixed model. We were not primarily interested in differences between experiments in microsaccade behavior, but including experiment as a factor in the model allowed us to test for interactions between experiment and precue type, our main variable of interest. We statistically analyzed microsaccade rate in the 500 ms before T1, the pre-target inhibition period during which the mean microsaccade rate was decreasing approximately linearly.

We used a two-stage procedure to assess the effect of precue type on microsaccade rate in this time window and determine significant clusters of time points. In the first stage, we tested whether the effect size at each time point was larger than expected by chance. For each time point (every 1 ms), we fit a linear mixed model to the mean microsaccade rate for each observer, with precue type and experiment as fixed-effects factors and observer as a random-effects factor. We used treatment contrasts. For all reported tests, the base conditions were “precue T1” for precue type and “Experiment 1” for experiment. To assess the significance of a difference between conditions (e.g. precue T1 vs. neutral) at the time point level, we compared the beta value estimated for that difference to the corresponding distribution of beta values from 1,000 permuted sets of the data, in which the precue type label was randomly shuffled for each observer. Time points at which the beta value of the real data fell in the upper or lower 2.5% of the permuted beta distribution were considered significant at the time point level.

In the second stage, to address the problem of multiple comparisons that arises at the timepoint level, we performed cluster-level tests (Maris and Oostenveld, 2007). These tests determined whether the total effect size across a cluster of individually significant time points was larger than expected by chance. For the real data and each permuted set, we defined clusters as contiguous time points with significant beta values and computed the sum of the beta values in each cluster. The maximum cluster sum from each permuted set was used to form a null distribution at the cluster level. A cluster in the real data was considered significant if its beta sum fell in the upper or lower 2.5% of the null distribution. The null distribution was used to calculate 2-tailed p-values for the clusters.

Finally, to control for any effects of experiment and assess interactions in each cluster, we performed an additional analysis of the cluster means. In each significant cluster we calculated the mean microsaccade rate for each precue type and observer, and we fit a linear mixed model to these cluster means. We then used a parametric bootstrap procedure for mixed models with 10,000 bootstraps to derive p-values.

#### Microsaccade timing analysis

To determine the effects of temporal attention on the precise timing of microsaccades around the time of the stimuli, we measured the onset latency on each trial of the last microsaccade before T1 onset (pre-T1 latency) and the first microsaccade after T1 onset (post-T1 latency) (Bonneh et al., 2015). We set the pre/post boundary at T1 onset, the start of the stimulus sequence. Few microsaccades occurred during the stimulus sequence. For the pre-T1 latency, we included microsaccades that occurred 1000-0 ms before T1, as these were the only microsaccades that could be affected by the precue. For the post-T1 latency, we included microsaccades that occurred from 0-2750 ms (the end of our microsaccade analysis window). Trials with no microsaccade in the relevant window were not included in the latency analysis.

To assess the effect of temporal attention on the timing of pre- and post-T1 microsaccades, we generated latency distributions for each precue type. To combine data across observers with different overall microsaccade rates and latencies, latencies from all trials for each observer were first z-scored regardless of precue type. As a summary metric, we calculated the median z-scored latency for each observer and precue type. To visualize the group latency distributions, the probability density of the z-scored latencies was estimated separately for each condition and observer by calculating the kernel density using 100 equally spaced points from −5 to 5. We plotted the mean and standard error of the density across observers and marked the median of each group distribution. *Statistics*. To evaluate the effects of precue type on mean pre- or post-T1 latency, we fit a linear mixed model to the median latency for each observer. We tested for main effects and interactions by approximating likelihood ratio tests to compare models with and without the effect of interest. We tested for pairwise differences between conditions using a parametric bootstrap procedure for mixed models with 10,000 bootstraps, based on the beta values from the linear mixed model.

#### Microsaccade timing vs. behavior analysis

We performed three types of analyses to assess the relation between microsaccade timing and behavioral performance. First, we tested for microsaccadic suppression (lower performance when a stimulus closely follows a microsaccade) by comparing trials in which a microsaccade occurred in the interval 0-100 ms before each target to trials in which no microsaccade occurred in that interval. For each target, we calculated accuracy on trials in which that target was probed by the response cue, separately for microsaccade and no microsaccade trials for that target. We used a linear mixed model and approximated likelihood ratio tests to compare microsaccade and no microsaccade trials for each target.

Second, we tested the relation between last pre-T1 and first post-T1 microsaccade latencies and behavior. We binned the trial period into 200 ms time intervals. For each latency bin and target, we calculated the change in accuracy with respect to a baseline, the mean accuracy across all trials in which that target was probed. We used a linear mixed model to compare the change in accuracy for each target and latency bin to zero (no change in accuracy when a microsaccade occurred at that latency). We corrected for multiple comparisons using a Bonferroni correction across all latency bins and targets.

Third, we assessed microsaccade-contingent behavioral tradeoffs between T1 and T2 at a higher temporal resolution (100 ms bins, 10 ms step size). T3 data was noisy at this resolution due to the smaller number of observers, so we focused on T1 vs. T2. We used the same analysis as just described to calculate the change in accuracy with respect to the baseline for each latency bin and target. We used a linear mixed model and approximated likelihood ratio tests to compare the values for T1 and T2 (i.e., did a microsaccade at a specific time change performance differentially for T1 and T2?). We then performed a cluster-corrected permutation test across time (see *Microsaccade rate analysis: Statistics*) to determine whether any time windows showed a significant difference between T1 and T2.

#### Code accessibility

Code for the behavioral experiments is available on GitHub at http://github.com/racheldenison/temporal-attention. Microsaccade analysis code is available at the following repositories: https://github.com/racheldenison/ta-microsaccades, https://github.com/racheldenison/ta-stats-R.

## Results

### Behavior

Thirty human observers judged the orientations of grating stimuli appearing in short sequences. Temporal attention was manipulated to different stimulus times using a precue (**Figure 1**). The precue could be informative, a single tone indicating the target likely to be probed with 75% validity, or uninformative (neutral).

Temporal precueing improved orientation judgment accuracy, as previously reported in Denison et al. (2017), Experiments 1-3. All datasets used for the microsaccade analysis, including replacement datasets, are combined here and replotted (**Figure 1d**). To quantify the behavioral effect of temporal attention in the combined data, we analyzed normalized accuracy. Accuracy was highest for valid trials, intermediate for neutral trials, and lowest for invalid trials, for both T1 and T2 (main effect of validity, T1: χ^2^(2) = 20.33, p = 3.8 × 10^-5^; T2: X^2^(2) = 13.54, p = 0.0011), with a performance increase from invalid to valid trials of 16% for T1 and 12% for T2. The improvement with attention was comparable for the two targets (no interaction between validity and target, X^3^(2) = 0.83, p = 0.66). This improvement did not depend on the experiment (no interactions between validity and experiment or among validity, target, and experiment, X^2^ < 3, p > 0.6). These behavioral data show that temporal attention was successfully manipulated in the current data set, which allowed us to ask how temporal attention affected microsaccades.

### Microsaccade detection

During the experiments, online eye tracking was used to detect and repeat trials with blinks or gaze position >1.5° from fixation. Therefore, completed trials did not contain blinks or large eye movements. Microsaccades <1° were detected offline with standard algorithms (Engbert and Kliegl, 2003) and followed the main sequence, with a mean correlation between amplitude and velocity across observers of 0.88 (SD 0.034).

### Microsaccade rate

The overall microsaccade rate exhibited expected dynamics across the trial period (**Figure 2a**). Mean microsaccade rate was ~1.8 Hz at the time of the precue tone, 1000 ms before T1. Following the tone, the rate dipped, rebounded, and returned to the baseline level over the course of 300 ms; these are characteristic dynamics following an auditory stimulus (Rolfs et al., 2008; Yuval-Greenberg and Deouell, 2011). Microsaccade rate then decreased approximately linearly during the 500 ms before T1 from a rate of ~1.7 Hz to a rate of ~0.2 Hz. We refer to this decrease as “pre-target inhibition.” The near complete inhibition we observed indicates a strong effect of stimulus timing expectations on microsaccades in our task. During the target presentations, the microsaccade rate remained near zero. It then rebounded 300-500 ms after T1 (“post-target rebound”). After an initial sharp rebound to ~0.7 Hz, microsaccade rate continued to increase slowly, reaching a value of ~1.5 Hz at 1000 ms after T1.

**Figure 2.**
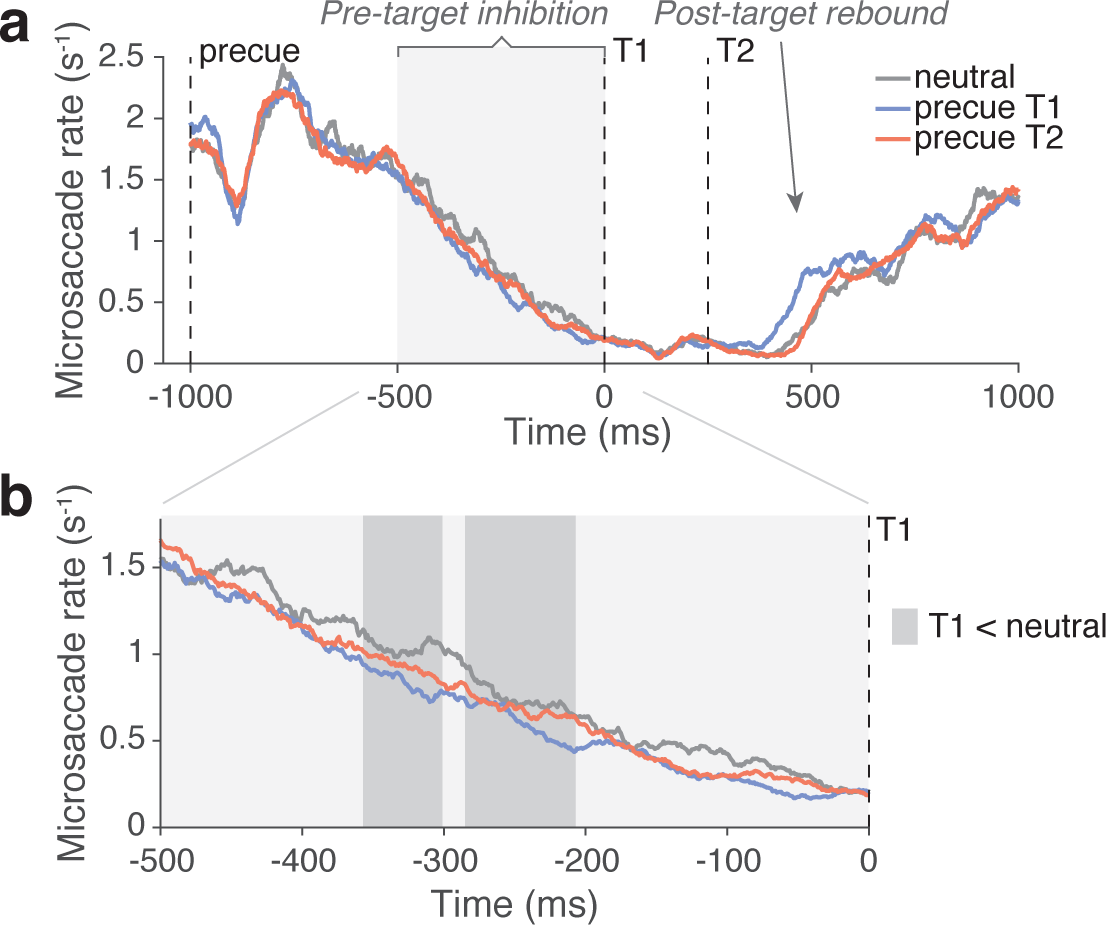
Microsaccade rate. **a)** Mean microsaccade rate across the trial. Data are combined across the common precue conditions of Experiments 1-3 (neutral, T1, T2, shown as separate colored lines). Dashed vertical lines show trial events. The response cue is not shown because its timing differs for 2-target and 3-target tasks. Light gray shading shows the pre-target inhibition period used for statistical analysis, and an arrow indicates the post-target rebound. **b)** Enlargement of pre-target inhibition period labeled in panel a. Dark gray shading shows significant cluster-corrected time windows, p < 0.05. n=30.

On top of these overall dynamics, the microsaccade rate was modulated by the precue. Microsaccade rate depended on precue type during the pre-target inhibition period (**Figure 2b**). Across this period, neutral trials tended to have the highest rate, precue T1 trials tended to have the lowest rate, and precue T2 trials had an intermediate rate. Microsaccade rate for neutral trials was significantly higher than the rate for precue T1 trials in two time windows (window 1: −357 to −301 ms, beta sum = 20.14, p = 0.01; window 2: −285 to −207 ms, beta sum = 28.02, p = 0.004; highlighted in dark gray in **Figure 2b**), as determined by a linear mixed model followed by cluster-corrected permutation tests. The mean values in these windows (window 1: neutral = 1.04, precue T1 = 0.84, precue T2 = 0.93; window 2: neutral = 0.75, precue T1 = 0.61, precue T2 = 0.69) showed no significant interactions between precue type and experiment (all absolute beta < 0.34, p > 0.05). Therefore, during the pre-target inhibition period leading up to T1, microsaccade rate was lower when T1 was precued than when the precue was uninformative.

The mean timeseries also showed an earlier post-target rebound when T1 was precued compared to when other precues were given (**Figure 2a**). However, because the rebound occurred at different times for different observers and experiments (rebounds were systematically later when there were three targets), it seemed inappropriate to statistically assess the mean microsaccade rate during the rebound period. Instead, to assess within-observer microsaccade timing shifts as a function of the precue, we quantified rebound timing, as well as the timing of pre-target inhibition.

### Microsaccade timing

To investigate the effect of temporal attention on precise microsaccade timing in the temporal vicinity of the target, we quantified the pre-target inhibition and post-target rebound timing. **Figure 3a** shows a raster of microsaccade onset times for an example observer. For each trial, the pre-target inhibition latency was defined as the onset latency of the last microsaccade before T1 (“last pre-T1 MS”), and the post-target rebound latency was defined as the onset latency of the first microsaccade after T1 (“first post-T1 MS”). This measure has been called “msRT,” as it is analogous to a reaction time measure (Bonneh et al., 2015). We used T1 as the reference time, because it is the start of the target sequence, so the pre-target period is free from oculomotor responses to the stimulus onsets. Few microsaccades were made during the target sequence, so most post-T1 microsaccades (93.3%) occurred after T2 as well.

**Figure 3.**
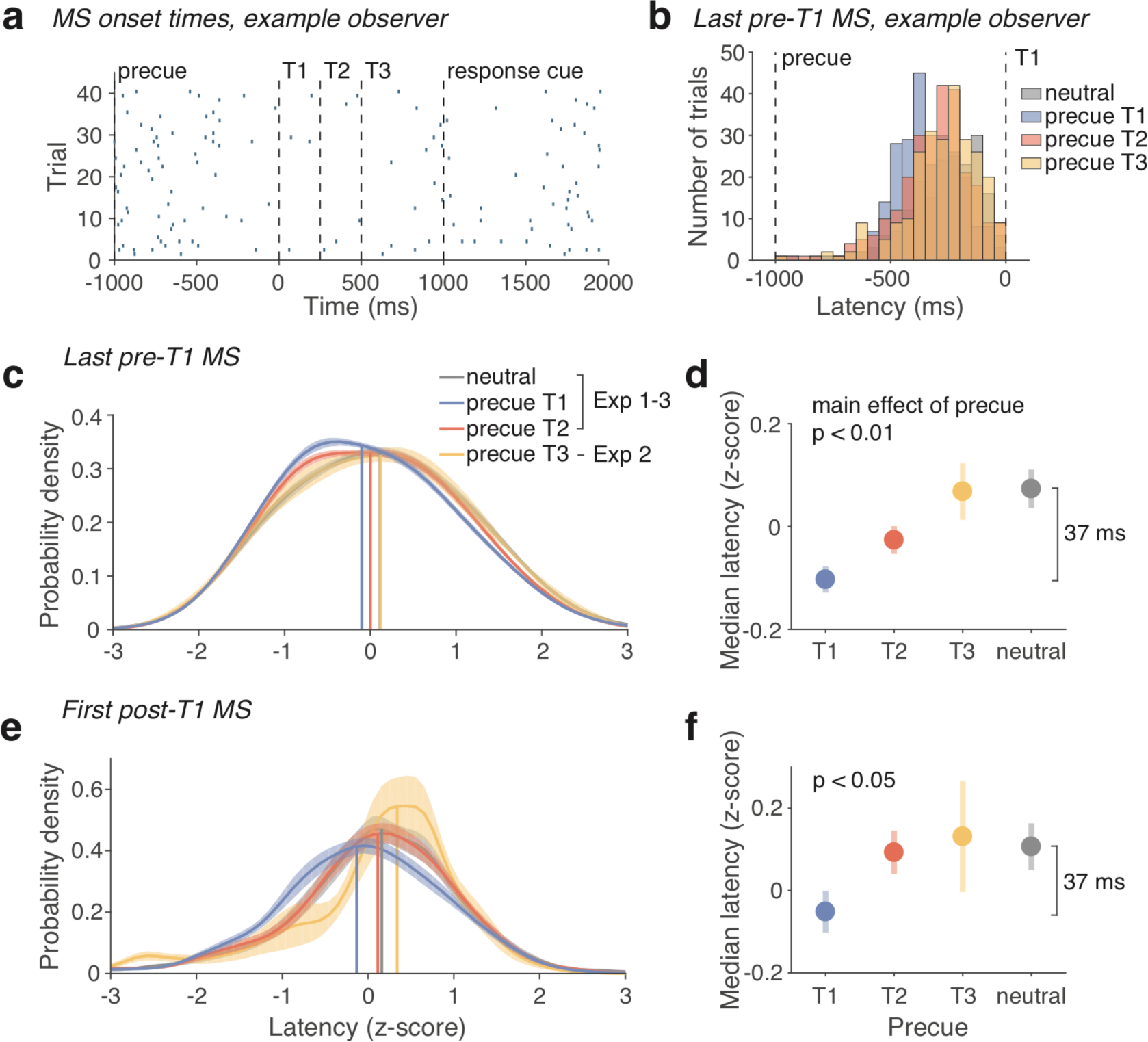
Microsaccade timing. **a)** Raster plot showing microsaccade (MS) onset times (blue ticks) in 40 precue T1 trials for an example observer from Experiment 2. Dashed vertical lines show trial events. For each trial, the latencies of the last pre-T1 MS and first post-T1 MS were recorded to quantify inhibition and rebound timing, respectively. **b)** Distribution across trials of inhibition (last pre-T1 MS) latencies for another example observer from Experiment 2. Precue conditions are shown in different colors. **c)** Inhibition latency distributions (last pre-T1 MS) for the group of observers. Latencies were z-scored to combine across observers with different overall timings and MS rates. Colored lines and shaded regions show mean and SEM of the estimated probability density for each precue condition. Vertical lines show the medians of the group distributions. **d)** Summary of group latencies. Markers and error bars show mean and SEM of each observer's median z-scored latency for each precue condition. The absolute magnitude of the latency difference between precue T1 and neutral conditions was 37 ms. **e)** and **f)** correspond to panels c and d but show the rebound (first post-T1 MS) latencies. Experiments 1-3, n=30; Experiment 2, n=9.

We evaluated the distributions of pre-T1 and post-T1 latencies across trials for each precue type. **Figure 3b** shows the distributions for one example observer. Latencies varied across observers. Median pre-T1 latencies ranged from −713 ms to −297.5 ms, and post-T1 latencies ranged from 316 ms to 1069 ms. Post-T1 latencies were also systematically later for Experiment2, which had three targets, compared to Experiments 1 and 3, because each target presentation inhibits microsaccades. Therefore, to combine data across observers and experiments, pre-T1 and post-T1 latencies were first z-scored for each observer, regardless of precue type.

Microsaccade timing just before and after T1 depended on the precue type. Latencies followed a systematic temporal progression: earliest for precue T1 trials, later for precue T2 trials, and latest for precue T3 trials. We found this same progression for pre-T1 inhibition latencies (**Figure 3c**, summarized in **Figure 3d**) and post-T1 rebound latencies (**Figure 3e**, summarized in **Figure 3f**). The timing distributions for neutral trials were relatively late, similar to precue T3 for pre-T1 inhibition latencies and to precue T2 for post-T1 rebound latencies (**Figure 3c-f**).

The effect of temporal attention on microsaccade timing seen in the full latency distributions was confirmed by statistical analysis of the median pre- and post-T1 latencies for each observer and precue type (**Figure 3d,f**). There was a main effect of precue type on the median pre-T1 latency (χ^2^(2) = 16.59, p = 0.00025) and no interaction with experiment (X^2^(4) = 2.41, p = 0.66). Pre-T1 latency was earlier for precue T1 than for neutral trials (beta = 0.18, p < 0.001). It was also earlier for precue T2 than for neutral trials (beta = 0.19, p = 0.012). In the 3-target experiment, pre-T1 latency was earlier for precue T1 than for precue T3 trials (beta = 0.21, p = 0.006), despite the reduced power in this smaller data set. There was also a main effect of precue type on the median post-T1 latency (X^2^(2) = 8.33, p = 0.016) and no interaction with experiment (X^2^(4) = 5.03, p = 0.28). Post-T1 latency was earlier for precue T1 trials than for neutral trials (beta = 0.16, p = 0.005) and precue T2 trials (beta = 0.14, p = 0.017). Precue T1 did not differ from precue T3 in the 3-target experiment (beta = 0.21, p = 0.096). In the full dataset, no other pairwise comparisons between precue types were significant for pre-T1 or post-T1 latencies (all absolute beta < 0.09, p > 0.05). We confirmed that the results were similar when we analyzed the median or mean latencies without z-scoring. In unnormalized units, the average shift of the median latency from precue T1 to neutral trials was 37 ms for both pre-T1 and post-T1 latencies (**Figure 3d,f**). The similarity of the effects of the precue type on inhibition and rebound latencies suggests that with temporal attention, the period of microsaccadic inhibition simply shifts depending on which target is most relevant.

To better understand the nature of the inhibition and rebound microsaccades, we assessed whether observers had any directional bias toward the target location in the lower right quadrant of the screen. We found no evidence for such a bias (**Figure 4**). Rather, these microsaccades tended to have the typical horizontal bias (Engbert and Kliegl, 2003; Tse et al., 2004; Hermens and Walker, 2010), with a slight additional upward and rightward skew. Post-T1 microsaccades tended to have more of an upward bias than pre-T1 microsaccades. We also analyzed the directions of the inhibition and rebound microsaccades as a function of their latency (in 200 ms time bins) and the type of precue. As in the combined data, no time bin or precue condition showed a bias toward the target location.

**Figure 4.**
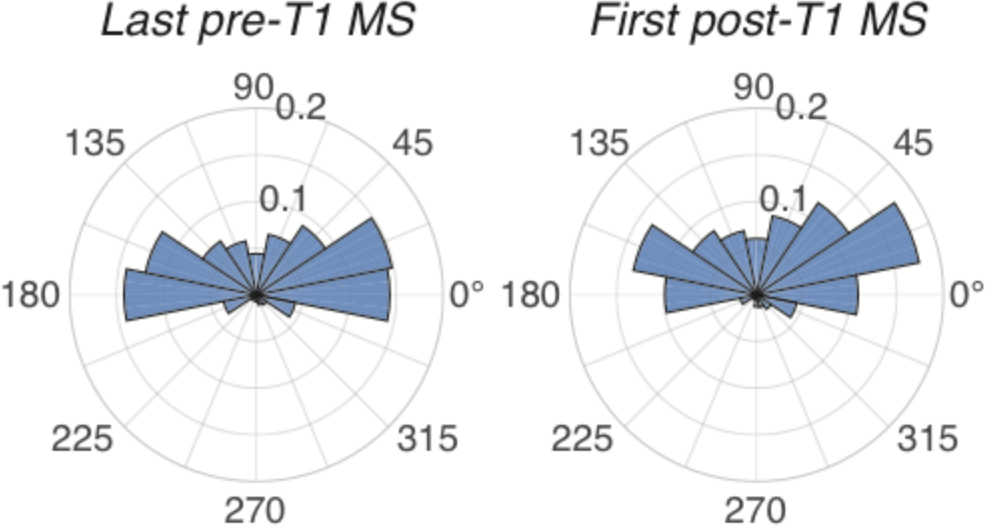
Directions of inhibition (last pre-T1) and rebound (first post-T1) microsaccades. Polar
histograms show the proportion of trials with microsaccades in each direction, out of the total number of
trials with pre-T1 / post-T1 microsaccades. Target stimuli were positioned at 315°.

### Relation between microsaccade timing and behavior

The effect of voluntary temporal attention on microsaccade timing predicts a functional relation between microsaccade timing and behavior. Such a relation has been documented in the form of microsaccadic suppression, or a reduction of behavioral performance when microsaccades occur just before a brief target (Zuber and Stark, 1966; Beeler, 1967; Hafed and Krauzlis, 2010; Hafed et al., 2011). In our data, consistent with these observations, accuracy in reporting the orientation of T1 tended to be lower when a microsaccade occurred 0 to 100 ms before T1 compared to when no microsaccade occurred in that time interval (X^2^(1) = 3.52, p = 0.061) (**Figure 5a**). This was not the case for T2 or T3 (X^2^(1)<1, p>0.3). Note that there were few trials with microsaccades in these time intervals, especially before T2 and T3 (**Figure 2a**), which could have reduced the quality of the accuracy estimates.

**Figure 5.**
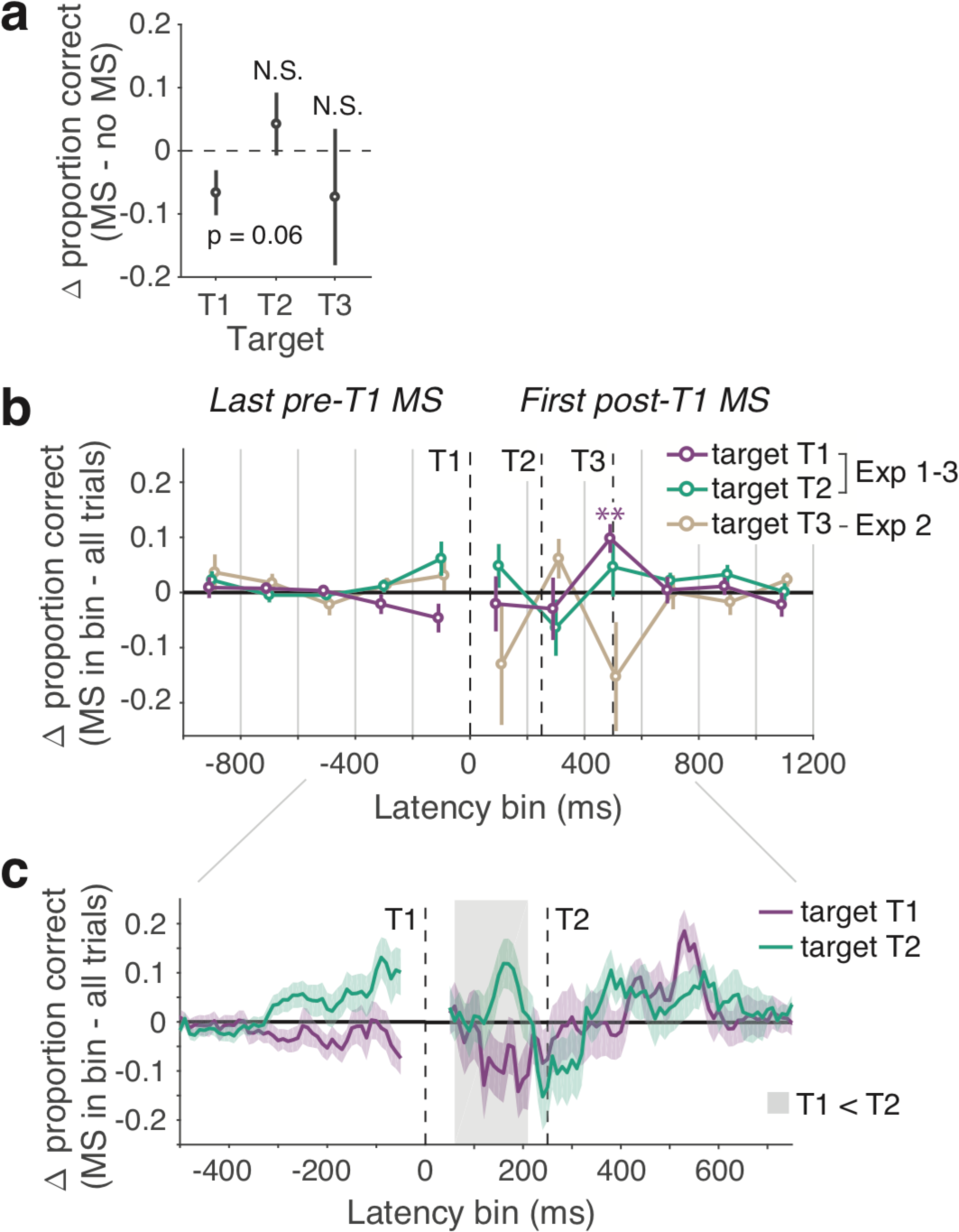
Relation between microsaccades and behavior. **a)** Test of behavioral microsaccadic suppression. Change in the accuracy of target report when a microsaccade occurred 0-100 ms before the target compared to when no microsaccade occurred in that interval. **b)** Effect of last pre-T1 and first post-T1 MS latency on behavior. Latencies are binned into 200 ms intervals (separated by gray vertical lines). Markers show change in accuracy when a MS occurred in a bin compared to mean accuracy across all trials for a given target. Mean and SEM are shown for each target (colored markers and lines). Dashed vertical lines show trial events. Experiments 1-3, n=30; Experiment 2, n=9. **p < 0.01. **c)** Same as the central portion of panel b but with higher temporal resolution (100 ms latency bins, 10 ms steps), to assess MS-driven behavioral tradeoffs between T1 and T2. (T3 not re-plotted at higher resolution because of lower reliability due to fewer observers.) Gray shaded region shows significant cluster-corrected time window for the difference between T1 and T2, p < 0.05. n=30.

We next assessed the relation between microsaccade timing and behavior specifically for the last pre-T1 and first post-T1 microsaccades used to quantify inhibition and rebound timing, respectively. Based on trial numbers, we binned the trial period into 200 ms time intervals. For each latency bin and target, we calculated the change in accuracy when a microsaccade occurred in that bin with respect to a baseline, the mean accuracy across all trials in which that target was probed (**Figure 5b**). Again consistent with microsaccadic suppression, microsaccades were associated with below average accuracy for a given target when they occurred in the same bin as that target, though these reductions did not reach significance. Interestingly, when the first post-T1 microsaccade occurred between 400 and 600 ms, orientation judgments for T1 were more likely to be correct (beta = 0.051, p = 0.002 following Bonferroni correction for multiple comparisons across all bins and targets). This timing corresponds to the post-target rebound timing evident in the microsaccade rate time series (**Figure 2a**).

We noticed that microsaccades in certain latency bins seemed to be associated with better performance for one target but worse performance for a different target. To evaluate the possibility that microsaccades at certain times contribute to performance tradeoffs between targets, we plotted the data for T1 and T2 only at a higher temporal resolution (100 ms bins, 10 ms step size) (**Figure 5c**). We then directly compared T1 and T2 microsaccade-related performance changes across time. A rebound microsaccade at 60-210 ms affected T1 and T2 performance differentially, improving performance for T2 but impairing it for T1 (beta sum = 2.54, p = 0.03). Note that T1 and T2 performance were not individually impaired or improved beyond chance levels; only the difference between T1 and T2 was significant. A similar difference between T1 and T2, though non-significant, was evident for microsaccades just before T1 (**Figure 5c**).

In summary, the timing of the microsaccades was behaviorally relevant in this task. We found microsaccadic suppression, enhanced behavioral performance for T1 when the rebound was ~500 ms after T1, and tradeoffs in performance between T1 and T2 contingent on microsaccade timing. These results confirm the relevance of attention-related microsaccade changes to visual sensitivity.

## Discussion

### Microsaccades reveal anticipatory mechanisms of temporal attention

When observers directed voluntary temporal attention, microsaccade rate decreased and microsaccade timing advanced, resulting in earlier microsaccadic inhibition in anticipation of the attended stimulus. Microsaccade rate decreased overall leading up to the predictable T1 onset, and it was lower in advance of the targets when the precue instructed observers to attend to T1 than when the precue was uninformative (neutral). The timing of complete microsaccadic inhibition before target onset also shifted systematically depending on the precue, with the earliest inhibition when T1 was precued. The timing of post-target rebound microsaccades shifted similarly, with the earliest rebound when T1 was precued. Thus, microsaccade dynamics both before and after the targets depended on which target time was instructed to be most relevant trial by trial, such that fixational stability increased around behaviorally relevant times.

Stabilizing fixation should benefit performance, given performance impairments when saccades or microsaccades occur during or just before brief targets (Ditchburn and Ginsborg, 1952; Zuber and Stark, 1966; Beeler, 1967; Herrington et al., 2009; Hafed and Krauzlis, 2010). Indeed, we confirmed that microsaccades occurring near targets affected behavior in our task. The visual system therefore stabilizes fixation not only based on predictable target timing (expectation) (Pastukhov and Braun, 2010; Hafed et al., 2011; Dankner et al., 2017; Amit et al., in press) but also based on task goals that change which stimulus time is most relevant from trial to trial (attention). This flexible adjustment of oculomotor behavior may be an overt mechanism of voluntary temporal attention.

These findings advance our understanding of the mechanisms underlying voluntary temporal attention. First, they provide a link between the cognitive process of prioritizing a specific moment in time and the activity of the oculomotor system. This link suggests that subcortical areas mediating eye movements receive temporal attention-related signals. The most likely pathway would involve top-down modulation of superior colliculus (SC) activity (Hafed et al., 2009), predicting changes in SC dynamics with temporal attention. Future studies could investigate interactions among SC; the frontal eye field, which projects to SC and contributes to the voluntary control of eye position (Martinez-Conde et al., 2013); and left intraparietal sulcus, which has been implicated in voluntary temporal attention (Coull and Nobre, 1998; Cotti et al., 2011; Davranche et al., 2011), to investigate the possibility that such a network contributes to the top-down control of microsaccade timing. Another candidate neural substrate is the striatal dopaminergic system, which has been suggested to mediate the effects of temporal expectations on oculomotor inhibition (Amit et al., in press).

The current findings also inform the debate on how early in time voluntary temporal attention affects visual processing, given that oculomotor changes have sensory consequences. Initial reports found late (>200 ms post-stimulus) effects of temporal precueing on stimulus-evoked neural responses (Miniussi et al., 1999; Griffin et al., 2002), along with pre-stimulus modulations of the contingent negative variation (CNV) of the EEG (Miniussi et al., 1999; Correa et al., 2006; Mento et al., 2015). These effects were attributed to cognitive and motor processes, such as preparing to respond. Subsequent studies found earlier effects on visual evoked responses (100 ms post-stimulus) (Correa et al., 2006; Anderson and Sheinberg, 2008), as well as a modulation of pre-stimulus alpha phase (Samaha et al., 2015). A separate line of research has shown that warning signals and hazard function manipulations can change visual cortical activity in anticipation of a target (Ghose and Maunsell, 2002; Müller-Gethmann et al., 2003; Cravo et al., 2011; Lima et al., 2011; Sharma et al., 2014; Snyder et al., 2016; van Ede et al., 2018). These manipulations inform observers about the probability that a target stimulus will appear at a given time. But they cannot dissociate expectation and attention, because the expected stimulus is always task relevant. Here, we dissociated these processes with a new task that manipulates temporal attention while controlling for expectation (Denison et al., 2017). Using microsaccades as a continuous physiological readout, we found clear evidence of preparatory processes associated with voluntary temporal attention up to 350 ms before stimulus onset.

### Temporal attention and expectation

Temporal attention and temporal expectation both contributed to microsaccade dynamics. Expectation was indicated by the inhibition of microsaccades, regardless of precue type, in the 500 ms leading up to the first target, such that the mean microsaccade rate was near zero when the sequence presentation began. The magnitude of the rate reduction was about 1.5 Hz. This is larger than the reductions of ~0.4 to 1 Hz found previously (Betta and Turatto, 2006; Pastukhov and Braun, 2010; Hafed et al., 2011; Fried et al., 2014; Dankner et al., 2017; Amit et al., in press). Individual observer variability could contribute to rate reduction differences across studies. Our observers received training on the task before the experiment, so their familiarity with the stimulus timing could have increased the expectation effects. It may also be that when stimulus timing is explicitly task relevant–as in our temporal attention task–the oculomotor system becomes more sensitive to stimulus timing overall. The temporal attention task we used requires temporal estimation (Grondin, 2010) of the 1000 ms interval between the precue and T1. Humans reproduce 1000 ms intervals with standard deviations of 200-350 ms (Lewis and Miall, 2009). Given this estimation uncertainty and the penalty of a microsaccade during a brief target, the optimal strategy would be to inhibit microsaccades early.

### A shift of microsaccadic inhibition?

Both rate and timing analyses suggested a simple shift of the pre-target inhibition and post-target rebound dynamics as a function of the temporal precue. In particular, the similarity of the pre- and post-T1 timing changes suggests a single, actively controlled inhibition process that shifts in time depending on which moment is most relevant. This account, however, requires further testing, and it does not perfectly predict some aspects of the current data. For example, post-T1 distributions were sharper and more skewed than pre-T1 distributions overall, likely due to the stimulus-driven component of the rebound. Indeed, whereas pre-target microsaccade dynamics are endogenously driven, post-target dynamics have both endogenous and stimulus-driven components, which could contribute to pre/post asymmetries. Based on the present data, we suggest a simple shift of inhibition timing as a parsimonious explanation for the effect of voluntary temporal attention on the endogenous component of microsaccadic dynamics.

Such an account raises the question: why would the system shift the timing of inhibition trial by trial rather than maximally sustain inhibition throughout the target presentation for all trials? One possible answer is that, in computational terms, there is a cost to maintaining perfect fixation. Observers can inhibit microsaccades voluntarily, for example when instructed (Steinman et al., 1967; Haddad and Steinman, 1973; Winterson and Collewijn, 1976), but doing so requires active control. The finding that a more difficult non-visual task during fixation is associated with fewer but larger microsaccades also demonstrates cognitive influences (Siegenthaler et al., 2014). Cognitive control over microsaccadic inhibition must interact with ongoing oculomotor dynamics. These dynamics can be described by a self-paced, stochastic process that generates saccades at semi-regular intervals (Amit et al., 2017). Both visual stimulus onsets (Engbert and Kliegl, 2003; Rolfs et al., 2008) and saccades (Nachmias, 1959; Beeler, 1965) produce saccadic inhibition, or a saccadic refractory period, which is followed by the next saccade (Otero-Millan et al., 2008; Amit et al., 2017). Such dynamics depend on the neural circuitry governing eye movement generation, which consists of multiple mutually inhibitory loops (Martinez-Conde et al., 2013; Otero-Millan et al., 2018). It may therefore be easier to shift periods of inhibition than to prolong them. Consistent with this idea is our finding that neutral trials showed shifted rather than prolonged inhibition dynamics–even though in neutral trials observers were instructed to attend to all targets equally. It will be interesting in future research to investigate the mechanisms underlying shifting vs. prolonging microsaccadic inhibition and how they relate to cognitive processes.

### Behavioral benefits and tradeoffs

So far we have focused on the detrimental effects of microsaccades on the perception of brief targets, which should promote the stabilization of fixation at relevant times. But our data also demonstrated that performance can be enhanced when microsaccades occur at specific times. In particular, we found that when observers made a rebound microsaccade 400-600 ms after T1, performance for reporting T1 was better than average. This improvement was specific to that time interval and to the T1 target. Relatedly, others have reported above-average performance when a microsaccade occurred 50-800 ms after target presentation when targets appeared in a rapid stream of stimuli (Pastukhov and Braun, 2010). The present data cannot determine whether the performance enhancement associated with a 500 ms rebound simply reflects successful inhibition during T1 or whether the rebound reflects some other neural dynamics associated with successful performance. The late timing of these rebound microsaccades, following intervening targets, suggests that the microsaccades are not changing the early-stage visual responses to T1 (which occur within ~200 ms). Therefore, this phenomenon likely differs from enhanced sensory processing for stimuli presented just before microsaccades (Chen et al., 2015). One study found better perception for stimuli occurring <100 ms after a microsaccade when microsaccades were directed toward vs. away from the stimulus (Yuval-Greenberg et al., 2014). However, it was not tested whether perception was better than if no microsaccade had occurred.

We also observed tradeoffs in behavioral performance across different targets as a function of microsaccade timing. Specifically, microsaccades between T1 and T2 (60-210 ms) were associated with differential changes to T1 and T2 performance, with relative impairments for T1 and improvements for T2. A similar, though less reliable, pattern was observed in the period leading up to T1. Microsaccade-contingent performance tradeoffs are interesting in light of the temporal attentional tradeoffs we have observed in behavior (Denison et al., 2017). Namely, voluntary temporal attention leads to both perceptual benefits for precued stimuli and perceptual costs for uncued stimuli, relative to performance following a neutral precue. Given the relation between temporal attention and microsaccades revealed in this study, these behavioral tradeoffs could be at least partially related to the timing of microsaccades.

## Acknowledgements

This research was supported by National Institutes of Health National Eye Institute R01 EY019693 and R01 EY016200 to MC, F32 EY025533 to RND, and T32 EY007136 to NYU, as well as by Binational United States / Israel National Science Foundation grant 2015201 to MC and SYG. We thank Stephanie Badde for statistical advice.

